# Exact mapping of Illumina blind spots in the *Mycobacterium tuberculosis* genome reveals platform-wide and workflow-specific biases

**DOI:** 10.1101/2020.03.11.987933

**Authors:** Samuel J. Modlin, Cassidy Robinhold, Christopher Morrissey, Scott N. Mitchell, Sarah M. Ramirez-Busby, Tal Shmaya, Faramarz Valafar

## Abstract

Whole genome sequencing (WGS) is fundamental to *M. tuberculosis* basic research and many clinical applications. Coverage across Illumina-sequenced *M. tuberculosis* genomes is known to vary with sequence context, but this bias is poorly characterized. Here, through a novel application of phylogenomics that distinguishes genuine coverage bias from deletions, we discern Illumina “blind spots” in the *M. tuberculosis* reference genome for seven sequencing workflows. We find blind spots to be widespread, affecting 529 genes, and provide their exact coordinates, enabling salvage of unaffected regions. Fifty-seven PE/PPE genes (the primary families assumed to exhibit Illumina bias) lack blind spots entirely, while remaining PE/PPE genes account for 55.1% of blind spots. Surprisingly, we find coverage bias persists in homopolymers as short as 6 bp, shorter tracts than previously reported. While GC-rich regions challenge all Illumina sequencing workflows, a modified Nextera library preparation that amplifies DNA with a high-fidelity polymerase markedly attenuates coverage bias in GC-rich and homopolymeric sequences, expanding the “Illumina-sequencable” genome. Through these findings, and by defining workflow-specific exclusion criteria, we spotlight effective strategies for handling bias in *M. tuberculosis* Illumina WGS. This empirical analysis framework may be used to systematically evaluate coverage bias in other species using existing sequencing data.

## INTRODUCTION

*Mycobacterium tuberculosis (M. tuberculosis)* is the leading cause of death from a single infectious agent, killing 1.5 million people globally in 2018 (1). Drug-resistance in *M. tuberculosis* is a major challenge for tuberculosis (TB) control and effective treatment (1). Today, whole genome sequencing (WGS) is the most commonly used tool for establishing new markers for TB surveillance and identifying candidates for molecular diagnostics as *M. tuberculosis* evolves (2, 3). Each sequencing technology has unique limitations, defined by both the sequencing instrument and library preparation (“sequencing workflow”). While WGS presents many opportunities for understanding and controlling TB, workflow-specific shortcomings are poorly understood. In this manuscript, we empirically evaluate depth of coverage across seven common Illumina workflows to describe workflow-specific coverage bias, which affects genome assembly and variant calling (4).

Illumina sequencing by synthesis (SBS) is by far the most commonly used WGS technology (5). Its library preparation includes a DNA amplification step, which significantly reduces the quantity of DNA required for sequencing. Illumina library preparation also allows many samples to be multiplexed in a single run, lowering sequencing costs substantially. These qualities make Illumina SBS desirable for many applications. However, some aspects of Illumina SBS limit its reliability for certain downstream analyses. Several genomic features cause biases that reduce coverage when sequenced on Illumina SBS technologies, particularly in segments of the genome where they are prevalent. The most well-characterized of these is GC content (4, 6, 7). GC bias originates primarily in library preparation, during amplification by polymerase chain reaction (PCR) (8). PCR is biased against amplifying GC- and AT-rich amplicons, which results in disproportionate read copy numbers (7, 8).

Repeat regions, homopolymers, and palindromes also reportedly drive coverage bias in short read sequencing (4, 7, 9–13). The inability of short reads to unambiguously span repeat regions prevents them from being mapped confidently to the genome, thereby reducing coverage depth (7). Homopolymers cause bias in some sequencing systems (4, 14), but Illumina states that homopolymers have virtually no effect on Illumina SBS (15). Palindromic sequences can form hairpin and stem-loop structures that can form during amplification and have been shown to impede sequencing (10, 16–18). Bias due to palindromes has been shown in sequencing-by-ligation technologies and long-read SBS, but reportedly does not introduce bias in Illumina sequencing (10, 17–19). Coverage bias due to these sequence attributes is influenced by two choices in the sequencing workflow: sequencing instrument and library preparation.

Many researchers are unaware that Illumina WGS data is affected by coverage bias, and take very low coverage to imply true deletions, ignoring coverage bias as a potential cause (20, 21). Others are aware of this bias, and handle it by excluding large regions of the genome that meet field-standard criteria with limited knowledge of which locations are affected (7, 22, 23). A common practice for handling Illumina sequencing bias in *M. tuberculosis* is to exclude all or part of the PE and PPE multigene families (23–26). These genes make up 10% of the coding capacity of the genome (27) and play roles in evading host immunity that are important, yet poorly understood (28). Indiscriminate exclusion of these families needlessly obscures valuable information from sites unaffected by coverage bias. Bias is not limited to PE/PPE genes, yet they are the primary excluded segments of the genome. Apart from PE/PPE genes (whose exclusion is often attributed to repetitive segments rather than GC-richness), researchers rarely address GC bias, despite its well-characterized contribution to coverage bias. This fracture in how the *M. tuberculosis* WGS community handles Illumina bias highlights a need for specific, empirically determined exclusion criteria. In an initial step towards this, Tyler and colleagues (6) studied the fidelity of Illumina-sequenced *Mycobacterium tuberculosis* genomes, and reported differential coverage bias between genomes prepared with Nextera and TruSeq library preps. They also reported that samples prepared with TruSeq resolved genomes into fewer contigs than Nextera and that certain regions, particularly GC-rich regions, could not be resolved with either library preparation method.

While these findings demonstrated differential coverage bias between library preps for *M. tuberculosis*, a systematic, single-base resolution analysis of positions in the *M. tuberculosis* genome that suffer from coverage bias is lacking. Here, we analyze coverage bias stratified across sequencing workflows to:

i. Provide lists of “blind spots” in the *M. tuberculosis* reference genome with consistent coverage bias.
ii. Characterize coverage bias for features of sequence composition known to be problematic.

We provide these deliverables for seven sequencing workflows separately, and in a pooled set. We find that blind spots distribute differently across the *M. tuberculosis* genome than accounted for by common practices in WGS analysis pipelines (7, 23–26) and overlap a variety of genes, including several implicated in drug resistance. Workflow-stratified analyses identify a modified Nextera library prep (29) as the least biased option and reveal distinct coverage biases across Illumina sequencing workflows, and the unexpected finding of coverage bias at homopolymers of a shorter length than previously thought. The provided blind spot lists (30) enable more informed interpretation of short-read sequencing data and improved WGS design for genome-wide association and phylogenomic studies.

## MATERIAL AND METHODS

### Sequence processing

First, we searched NCBI (National Center for Biotechnology Information) *Mycobacterium tuberculosis* genomes uploaded between January 1^st^, 2016 and March 24^th^ 2019 and sequenced on an Illumina instrument, producing 5,131 candidate genomes. Genomes without reported library prep, or prepared with a library prep other than a traditional or modified Illumina library were excluded. Only genomes from sequencing instrument and library prep combinations (sequencing workflows) with ≥20 total genomes were included. Candidate SRA IDs were pulled using sratoolkit’s (31) --prefetch function and raw fastq files were obtained from NCBI using sratools --fastq-dump. After each file was downloaded, genomes underwent reference-based assembly using a parallelized in-house pipeline: First, the downloaded raw reads were trimmed to eliminate adapter sequences and remove low quality ends using Trimmomatic (32). After removing artifacts, reads were aligned to the H37Rv reference genome (NC_000962.3) using bowtie2 (33). The samtools (34) package was then used to sort, index, and produce an mpileup file. Assembled genomes with low average coverage were then excluded, leaving 1,547 genomes. A custom python script was then implemented to find positions that met our low coverage criteria for each genome. Finally, VarScan2 (35) was used to identify variants for building phylogenies.

### Identifying low coverage positions

Sequencing depth, or “coverage” refers to the number of reads that map to a position during alignment, and average coverage is the mean of coverage across all positions in the genome. Instead of an absolute coverage threshold, we defined “low coverage” using a relative threshold specific to each genome. We express relative coverage of each position (*Di*) as the ratio of the detected coverage at that position (*d_i_*) to the average coverage (*μC_i_*) in the genome (Equation 1). We sought to determine which bases in each genome belonged to the set of low coverage positions (*K_LC_*) for that genome. A given position *k* belonged to *K_LC_* when its relative coverage was ≤ 0.1 (Equation 2).

### Phylogenetic filtering

When mapping reads to a reference, positions in the reference genome that are absent in the clinical genomes (true deletions) would be considered “low-coverage”. To account for this, we implemented a phylogenomic filtering step to identify and exclude true deletions from consideration as low-coverage positions on a genome-specific basis. First, a maximum likelihood phylogeny was created using RAxML (36) version 8.1.1 with a general time reversible model of evolution and 100 bootstrap replicates on a concatenation of 70,057 single-nucleotide polymorphisms (SNPs), gathered from each genome’s VCF file, which were generated using Varscan2 (from our reference-based assembly pipeline). *Mycobacterium bovis* and *Mycobacterium canetti* were used as outgroups. The tree was visualized and manipulated using the iTOL (37, 38) webtool.

Next, we found which genomes shared each low coverage position. For each position, the set of genomes with low coverage at the position were checked for monophyly on the phylogenetic tree, using the ete3 (39) python package, run on the newick file generated by RAxML (40). Each position with all low coverage members belonging to a monophyletic group of a size larger than the minimum number required to qualify as a blind spot (n = 5, calculated according to Equation 5) was considered a true deletion, rather than a potential blind spot. True deletions identified through these steps were removed from downstream analyses. Following filtering, remaining low coverage positions (Figure 1) were screened for polyphyletic groups containing monophyletic subgroups at least as large as the threshold for counting as blind spot (n = 5; Equation 5). In such cases, the genomes comprising these monophyletic subgroups were excluded from *G*, the total number of genomes containing the position of interest, when determining blind spot classification (Equation 5).

**Figure 1.**
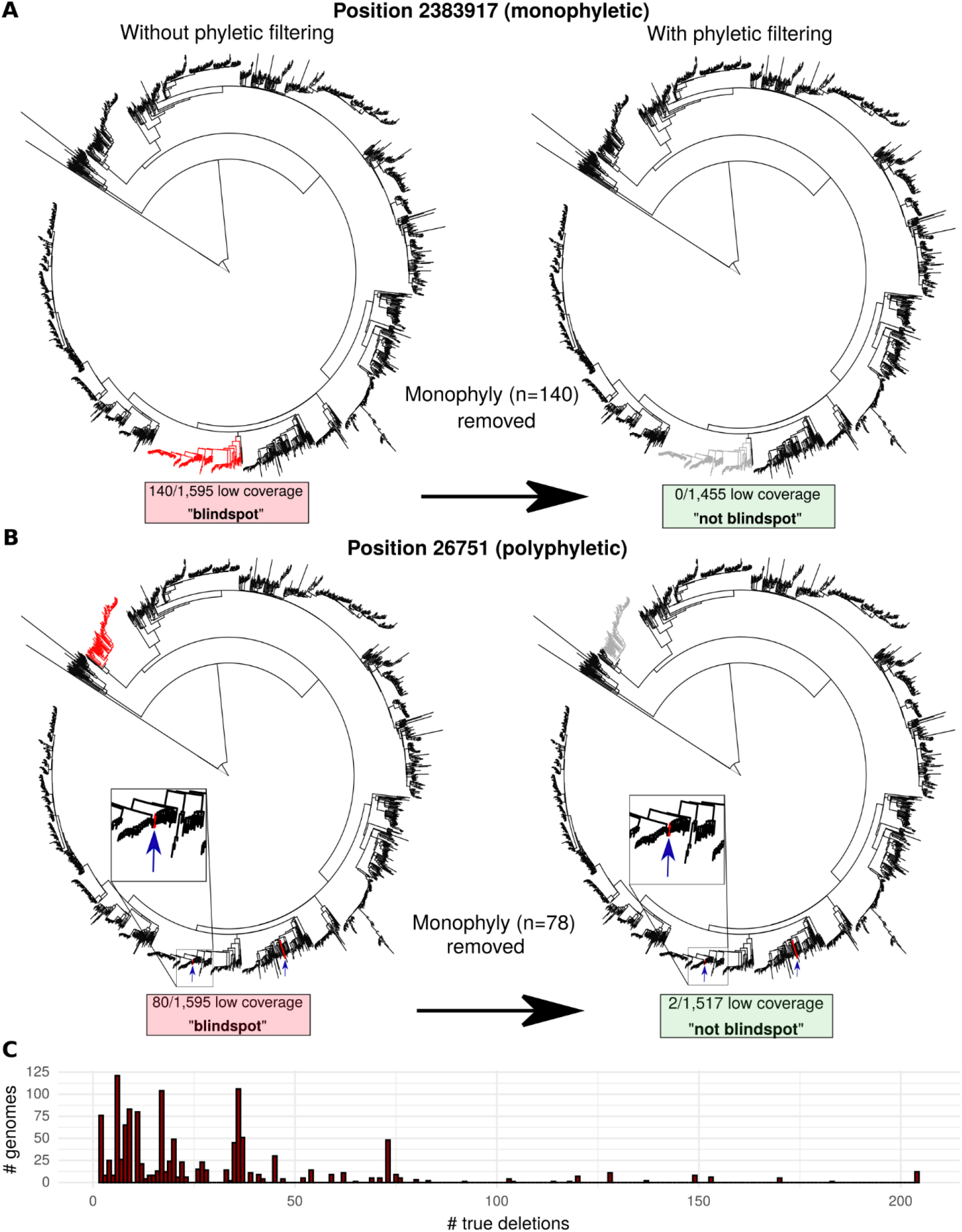
Phylogenetic filtering of true deletions reduces blind spot false discovery rate. True deletions can confound naïve coverage-based analysis. Our phylogenetic filtering method is depicted, showing positions where low coverage positions (red) were frequent enough to meet blind spot criteria when called naïvely. Example positions with (**A**) monophyletic and (**B**) polyphyletic distributions of genomes with low-coverage (<10% of mean genome coverage) are depicted prior (left) and following (right) phylogenetic filtering. (**C**) distribution of genomes harboring number of true deletions identified by phylogenetic filtering. The high number of genomes with identical numbers of true deletions suggests phylogenetic filtering captured clonal expansions harboring deletions that would have otherwise inflated the blind spot count.

### Classifying blind spots

We took a probabilistic approach to determine the threshold for how many genomes a low coverage position had to occur in to be considered a “blind spot.” In a genome, each position, *k*, was considered to have “low coverage” if the coverage at position *k* was less than *d_i_* (Equation 2). Each of the positions meeting this criterion were then included in set of positions with low coverage (*K_LC_*) for the genome. Positions that were true deletions in monophyletic groups and monophyletic subsets in polyphyletic groups were then excluded from *K_LC_* using the phylogenetic approach described above. Given the size of the set of positions included in *K_LC_*, we calculated the probability (*E*) that, by chance, a given position *k* belonged to *K_LC_* across genomes (Equation 3). Therefore, *E^n^* is the probability that a given position *k* has low coverage in *n* genomes, by chance (Equation 4). This probability was calculated separately for each instrument/library prep workflow.

Next, we calculated the probability (*P_LC_*) that position *k* had low coverage in *n* of *G* total genomes in the workflow by random chance (Equation 5). The value of *G* was specific to each position because a base with low coverage could have been a true deletion in some genomes and a potential blind spot in others (Figure 1B). In other words, for a given position, *G* excluded the number of genomes in which the position was truly deleted (determined using the monophyletic subsets in polyphyletic groups).

To determine the acceptable false positive rate (*F_p_*), we first set a threshold for the number of false positives we considered “acceptable” to include. We sought to capture a minimal set of blind spots and be conservative in the number of false positives included. With this objective in mind, we accepted only 0.1 false positives in our set of classified blind spots. A false positive rate of 6×10^−7^ yields 0.1 false blind spots, therefore we set (*F_p_*) to this value. Using this acceptable false positive rate (6×10^−7^) and our known genome size (4,411,532), we defined our blind spot detection threshold (*F*) as a function of the median (across genomes in each sequencing workflow) number of low coverage positions (*K_LC_*) that were not deleted (Equation 6).

For each position that had low coverage in a genome, *P_LC_* was calculated and compared to the blind spot detection threshold *F* to determine if the position had low coverage in more genomes than we would expect by chance. If the observed probability of a position appearing as low coverage in *n* genomes by chance out of *G* genomes in a workflow was lower than our detection threshold, we included that position in our final set of blind spots, *B_s_* (Equation 7).

### Annotating sequence attributes

#### Homopolymers

Homopolymers were defined as any sequence of consecutive (n > 1) identical bases (Guanine, Cytosine, Thymine, or Adenine). Homopolymers were retrieved from the H37Rv genome (NC_000962.3) with a custom shell script.

#### Repeats

To identify repeat regions, we used the Tandem Repeat Finder open source software (TRF) (41). TRF uses Smith-Waterman alignment to detect repeats, and filters candidate repeats based on alignment score values the user inputs.

#### GC content

Prior work has demonstrated that the primary driver of GC bias is the GC content (guanines + cytosines)/ total bases) of fragments during PCR amplification (42). For a given base, the probability that its flanking bases will co-occupy the same fragment diminish as a function of the distance between them. However, relatively distant bases (up to the length of the fragment) will sometimes contribute to GC content. To capture bases with extreme GC content in fragments at multiple scales that might contribute to GC bias, we independently considered GC content in variably-sized windows, calculating GC content around each base of the H37Rv genome for each window size (50 bp and between 100-1000 bp, in 100 bp intervals).

#### Palindromes

We identified palindromes in the H37Rv genome using EMBOSS (43) suite’s palindrome software. This component of the EMBOSS software package scans the genome for inverted matches, and filters based on user-defined match/mismatch and gap requirements. We included palindromes with stem length ≥ 7 (44) and allowed for mismatches and/or gaps based on EMBOSS’s recommended settings to capture a wide range of candidate palindromes.

### Defining thresholds to classify sequence attributes

We took an iterative, empirical approach to determine thresholds for the following attributes: homopolymer length, repeat length and period size, and “extreme” GC content. Within each iteration, sequencing attributes were binned on their criteria (e.g. *length* of homopolymer) and examined bin-wise for deviation from the “unexplained” blind spot fraction (blind spots in positions qualifying for none of the attributes). The bases classified as one or more of the other attributes in the previous iteration were excluded from this set of unexplained positions. Thresholds for each criterion were set at the first (i.e. least extreme) bin/category where blind spots were significantly more prevalent than blind spots in unexplained sequences (Two-sided Fisher’s exact, 2.5^th^ quantile of odds ratio > 2) and increased monotonically thereafter. We iterated through this process until thresholds for all three attributes stabilized. All statistical tests for determining thresholds were implemented in R. For the first iteration, homopolymers were binned by length and the fraction of blind spots in each bin were compared to the fraction of blind spots in all other positions in the genome. The minimum homopolymer length that was significantly enriched for blind spots was set as the threshold. The same process was performed on tandem repeat lengths, binned in intervals of 2. Shorter repeats that did not meet the threshold for length were then separated on period size and investigated for enrichment of blind spots, though none were significantly enriched.

The threshold for “extreme GC content” was determined empirically and calculated separately for each window size. GC content was calculated for the number of bases in the window, half on each side flanking the base of interest. GC content was then binned in 2% intervals for each window size.

In each bin, the proportion of blind spots was calculated (# blind spots matching criteria/total positions in genome matching criteria) and compared to the proportion of blind spots in sites “unexplained” by homopolymers and repeats. While all bins were investigated for bias (high and low GC content), only GC-rich bins had significantly disproportionate blind spot fractions. We therefore set thresholds only for high GC content for each window size. Bases were classified as considered “GC-rich” if their GC content exceeded thresholds for any of the windows.

### Instrument and library preparation error profiles

We identified blind spots separately for each combination (n=7) of instrument and library prep (sequencing workflow). The instruments included were NextSeq 500, MiSeq, HiSeq 2000, and HiSeq2500. Library preparations included were TruSeq, Shotgun Nextera, Nextera XT, and modified Nextera. We defined two sets of blind spots to be used for different purposes throughout the analysis: 1) A “pooled” set (n=1,547 genomes) and 2) a “comparison” set (n=175 genomes, 25 per workflow). After using the blind spots classified separately within each workflow using all available genomes (Supplementary Table S1), the union of these workflow-specific sets were pooled into a single “pooled set” (Supplementary Table S2) to capture additional blind spots. Since the blind spot criteria is conservative, favoring false negatives over true positives (Equation 6), evaluating additional genomes is likely to increase true positives with negligible increase in false positives. However, blind spot classification depends on the number of genomes considered, thereby adding the confounding effect of sample size when comparing the number of blind spots in each workflow. To enable fair comparison between workflows, we also determined the “comparison” set of blind spots by classifying blind spots using the same number of genomes within each workflow. The workflow with the smallest sample size (n=25) was the NextSeq 500 instrument paired with TruSeq library preparation. Genome-wide average coverage was then used to select 25 genomes from the other 6 workflows with the most similar average coverage. Blind spots were classified using these curated sets of genomes. This comparison set was used for the stratified parts of our analysis when contrasting the factors that challenge to each sequencing workflow.

### Statistical tests

#### Comparisons between Sequencing instruments, library preparation methods and their combinations

When comparing blind spots between sequencing workflows (combinations of instrument and library prep), statistical tests were chosen according to the number of groups being compared, whether they distributed normally, and whether they had similar variance. When more than two workflows were compared, ANOVA was used to compare all that distributed normally and had variances within the four-fold of one another (the “rule-of-thumb” for approximately equal variance of greatest variance must be no more than four times the smallest), and post-hoc Tukey tests were performed pairwise to estimate differences between means. Comparisons between non-normally distributed combinations were done pairwise using Wilcoxon signed rank sum test. Comparisons between the non-bootstrapped sequencing workflow were performed pairwise using one sample t-tests. Pairwise comparisons between normally distributing combinations of similar variance were evaluated with paired t-tests. All statistical tests were implemented in R (45).

### Annotating blind spots

Overlap between “pooled” blind spots and annotated coding regions in the H37Rv reference genome (NC_000962.3), downloaded from MycoBrowser (https://mycobrowser.epfl.ch/) was used to determine genes affected by blind spots. Genes containing blind spots were grouped by gene family and the fraction of bases in each family that were blind spots was calculated, along with the fraction of blind spots that were bases in genes of each family. To find genes implicated in drug resistance, we joined the gene-based blind spot list with a curated list of resistance-implicated genes from recent publications (Supplementary Table S3).

## RESULTS

The objective of this work was to precisely describe coverage bias of common Illumina sequencing workflows for *M. tuberculosis* WGS and translate them into actionable knowledge for researchers designing or interpreting the results of *M. tuberculosis* WGS studies. We took a phylogeny-aware, probabilistic approach to classify blind spots and stratified our analysis by sequencing instrument and library prep (sequencing workflow). We defined “low coverage” relative to each genome’s mean overall coverage (<10%), rather than an absolute threshold (e.g. <5 reads), mitigating the confounding effect of mean sequencing depth when applying an absolute threshold (Figure 2B). By analyzing coverage bias stratified across sequencing instrument and library preparation technique, we contrast how problematic sequence attributes challenge different sequencing workflows. We applied this approach to 1,547 recently sequenced genomes (Supplementary Table S1) across popular Illumina sequencing workflows to identify Illumina “blind spots” in the genome (30) that are systematically under-represented due to coverage bias.

**Figure 2.**
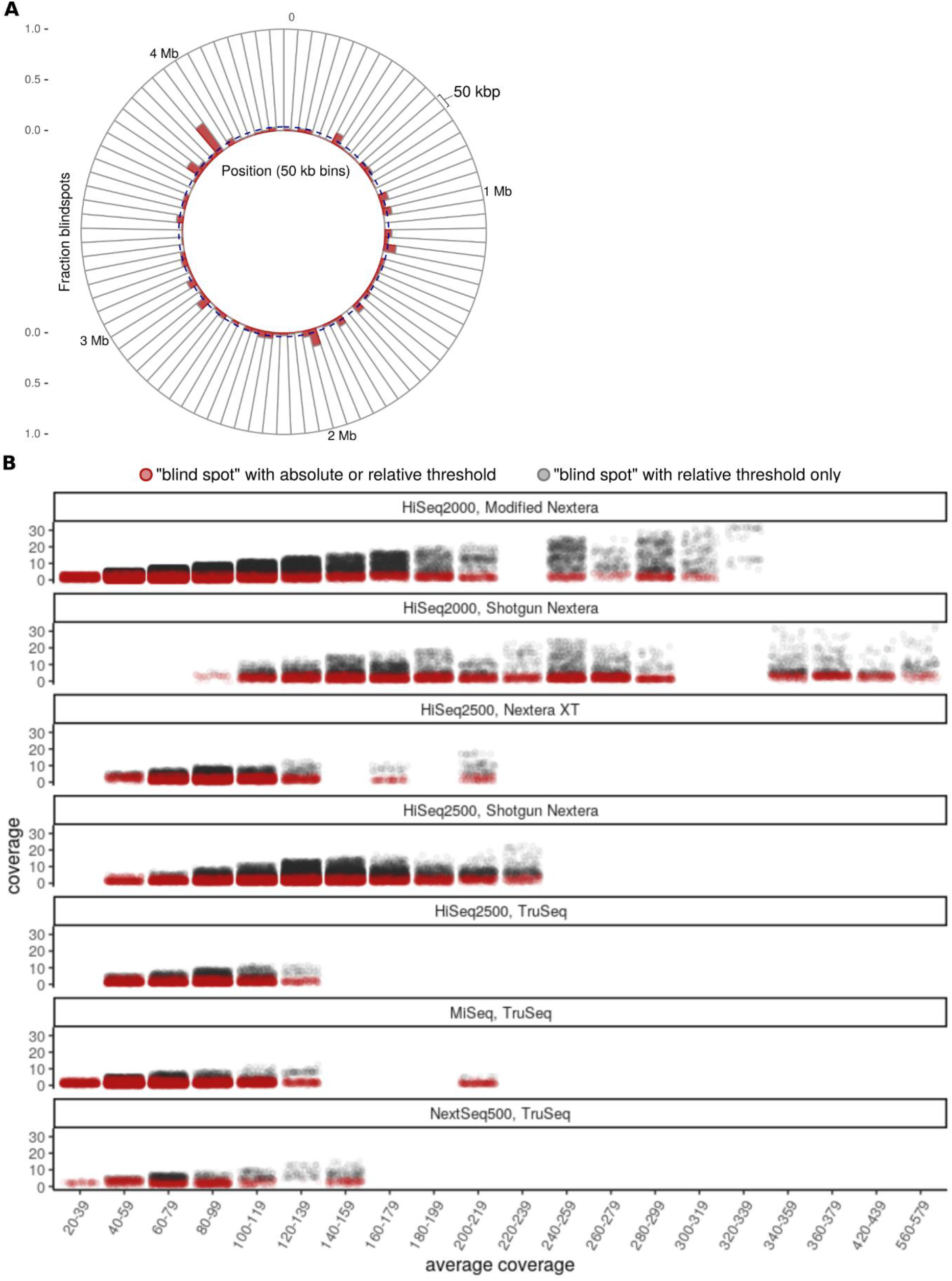
Illumina blind spots in *Mycobacterium tuberculosis* whole-genome sequencing. **(A)** Distribution of blind spots from the “pooled” set (the union across all seven sequencing workflows) across the genome of *M. tuberculosis* virulent type strain H37Rv. The H37Rv genome is binned into 50 kb segments and the fraction of blind spots (red) is shown for each bin. The dashed blue line represents the baseline fraction of blind spots across the genome (3.6%) highlighting areas with disproportionate levels of blind spots. (**B**) Comparison of blind spot classification only when using our relative threshold (gray) versus when using either our relative threshold or the previously used absolute (red) threshold (coverage < 5) (6) for classifying a position as having “low coverage”. Each position in the “pooled” set of blind spots is plotted according to its coverage (y-axis) and the average coverage (mean, x-axis) across the genome. Average coverage is binned (bin width = 20) and jittered within each bin, with each point rendered at 0.05 opacity to visualize density. Within any given sequencing workflow, the fraction of blind spots that are considered “low coverage” according to the relative threshold, but not the absolute threshold increases in step with average coverage. Therefore, when defining low coverage using an absolute threshold, genomes with higher average coverage appear to have fewer blind spots, whether or not this is truly the case.

### Phylogenetic filtering removes deletions masquerading as blind spots

If one assumes low coverage invariably indicates coverage bias during sequencing, true deletions would be included spuriously as blind spots. To remove such confounding phylogenetic events from our analysis, we filtered out positions where low coverage was localized to one portion of the phylogeny (monophyly) in more genomes than expected by chance. This filtered out positions both from monophyletic groups (Figure 1A) of genomes and monophyletic subsets (≥5 genomes) within polyphyletic groups of genomes (Figure 1B), removing true deletions without obscuring true blind spots. True deletion frequency (mean=23) distributed irregularly among genomes (Figure 1C), consistent with the clonal nature (46) of *M. tuberculosis* evolution. Following phylogenetic filtering, positions were classified as “blind spots” if they occurred in enough genomes within a sequencing workflow to meet our probability-based threshold (Equation 7).

### Catalogue of blind spots in the *M. tuberculosis* genome

We defined two sets of blind spots: A “pooled” set (n=1,547 genomes) and a “comparison” set (25 per each of the seven workflows, n=175 genomes). The pooled set is the union of blind spots across workflows, where blind spots are calculated separately for each workflow using all available genomes. This maximizes capture of true positive blind spots but is weighted unequally across sequencing workflows (due to inequity in genomes per workflow, Supplementary Table S4). This set includes blind spots present when using any of the seven workflows, which is useful for researchers who are using data from multiple workflows or workflows not included in this study. This pooled set comprises 3.6% (159,659 positions) of the H37Rv reference genome (Supplementary Table S2), scattered across the genome (Figure 2) in 5,888 regions. Only 1.1% of blind spots appear as single positions, while the majority appear in consecutive positions, forming clusters (mean length=27 bp, range 1-1,816 bp), consistent with the idea that coverage bias is driven by sequence context.

The comparison set of blind spots was created using the same number of genomes (n=25) from each sequencing workflow, accounting for the positive relationship between number of genomes in a group and number of blind spots classified in the group. In this comparison set (Supplementary Table S5), 8,379 (8%) blind spots were shared among all seven workflows, while most sites only have coverage bias for a subset of workflows. Many blind spots were specific to only one workflow (uniquely present) or were present in all other workflows but one (uniquely absent) (Table 1). Notably, 10,519 of the blind spots in the comparison set are uniquely absent among genomes sequenced with the HiSeq2000/modified Nextera workflow, more than all blind spots uniquely absent in the other six workflows combined. The intersection of blind spots between the remaining six workflows would increase from 8,379 to 18,898 if HiSeq2000/modified Nextera were not considered, a more than two-fold increase. This workflow is able to resolve more than half of the blind spots present among all other workflows considered, suggesting it mitigates bias in regions that challenge other sequencing workflows.

**Table 1.** Number of blind spots uniquely present or absent in each sequencing workflow.

| Instrument | Library Prep | Uniquely Present | Uniquely Absent |
| --- | --- | --- | --- |
| NextSeq 500 | TruSeq | 5359 | 3256 |
| MiSeq | TruSeq | 5726 | 1337 |
| HiSeq 2500 | TruSeq | 2590 | 451 |
| HiSeq 2000 | modified Nextera | 3314 | 10519 |
| HiSeq 2000 | Shotgun Nextera | 1869 | 158 |
| HiSeq 2500 | Nextera XT | 5387 | 2302 |
| HiSeq 2500 | Shotgun Nextera | 612 | 111 |

Tyler et al. reported 124 and 195 “ultra-low coverage” (ULC) hotspots in genomes prepared with TruSeq and Nextera libraries (6), respectively. These ULC regions were defined by arbitrary absolute low coverage (<5x) and length (>10 base pairs) thresholds. While we used a different version of the reference genome for assembly and different methods for identifying coverage bias, our blind spots capture a majority of their ULC regions (83% for TruSeq and 63% for Nextera). Our approach captured more coverage-biased sites, but the overlap between blind spots and Tyler and colleagues’ ULC positions show congruent results despite methodological differences, attesting to the reproducibility of coverage bias analyses.

### Sequencing workflow affects prevalence of blind spots in Illumina WGS studies

To compare blind spots between workflows we evaluated the number of blind spots in 100 bootstraps of 25 genomes (the minimum genomes in any given workflow, Supplementary Table S6). Our methods for estimating the expected difference in number of blind spots between sequencing workflows differed according to how the number of blind spots distributed among the bootstraps (Methods and Materials). However, when asking whether a given sequencing workflow produces *significantly* more/fewer blind spots than another workflow, we employed methods to capture what difference would be meaningful to researchers for WGS experimental design. When designing sequencing experiments, researchers often operate under significant financial or logistical constraints. When opting for a sequencing workflow that requires a more expensive library preparation or use of an instrument outside of their institution or trusted collaborators, they likely want to be confident that their sequencing experiments deliver fewer blind spots every time, or nearly so. Taking this into consideration when comparing sequencing workflows, we qualified the number of blind spots between workflows as “significantly” different only when the number of blind spots differ >99% of the time, rather than considering an arbitrarily small difference between means as significantly different. We use the relationship described by Payton & colleagues (47) to estimate this from the overlap of the range in 2.5^th^-97.5^th^ quantiles of two distributions under comparison.

Before comparing sequencing workflows, we examined the relationship between mean coverage among genomes in bootstraps and blind spots. While 3/6 sequencing workflows correlated significantly (*P* < 0.05, 2 negative, 1 positive) yet modestly with coverage (−0.27 < R < 0.29), the coverage-blind spot relationship does not appear to bias our conclusions regarding differences between workflows (Figure 3A).

**Figure 3.**
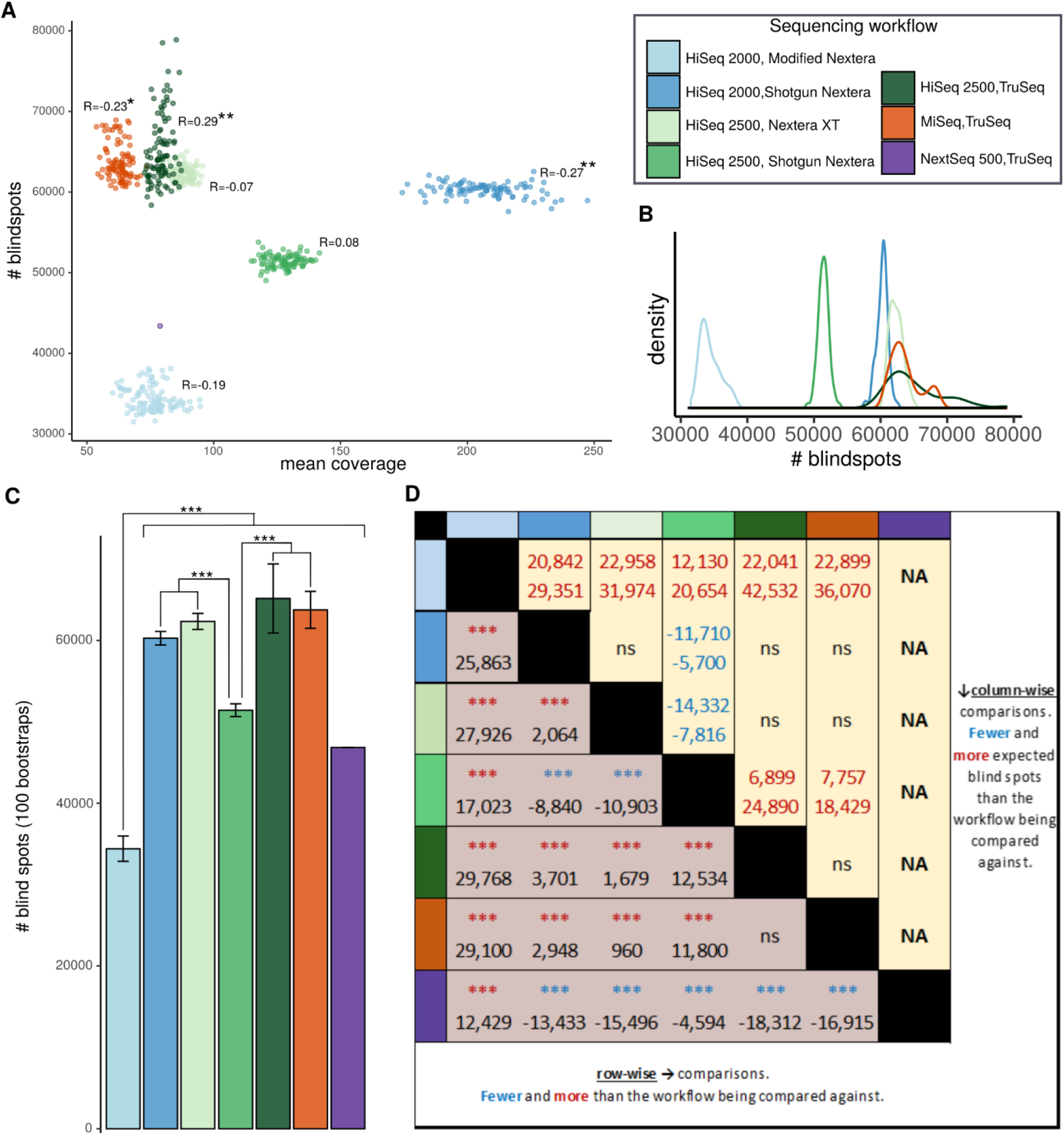
Blindspot prevalence across all instruments, library preps, and their combinations. (**A**) Scatterplot and correlations between the number of blind spots (y-axis) and mean coverage (x-axis) among 25 genomes for each bootstrap (n=100 per Instrument/Library Prep workflow). Correlation coefficients are displayed for each workflow \**P*<0.05, \*\**P*<0.01. (**B**) Distribution of the number of blind spots across bootstraps for each workflow. **(C)** The number of blind spots (y-axis) across the seven sequencing workflows (x-axis) (n=175, 25 genomes for each instrument/library prep workflow). Error bars represent ±1 standard deviation from the mean number of blind spots across bootstraps. ***non-overlapping 95% confidence interval. (**D**) Pairwise comparison between sequencing workflows of estimated mean blind spots (row-wise, bottom left) and expected difference in number of blind spots (99% interval, inferred by 95% confidence interval boundaries as previously described (47); column-wise, top right). Statistical tests were chosen according to distribution and equality of variance. Normal distributions of equivalent variance were compared with Tukey multiple comparisons of means; comparisons involving one or more sequencing workflows with non-normal distributions were compared with Wilcoxon rank sum test; one sample t-tests were used to compare the blind spots in the single NextSeq 500/TruSeq set to the mean blind spots across bootstraps in other workflows. \**P* < .01, \*\**P* < .001, \*\*\**P* < 1×10^−5^.

#### Coverage bias between instruments

We investigated the number of blind spots between libraries prepared with the same kit but sequenced on different instruments, focusing on evaluating differences in the expected number of blind spots when sequenced on each workflow (Figure 3D). We first contrasted blind spots between samples prepared with TruSeq, allowing us to compare HiSeq 2500, MiSeq, and NextSeq 500. Among these three instruments, NextSeq had the fewest mean blind spots (*P* < 2.2*10^−16^, one-sample t-test), while HiSeq2500 had modestly fewer blind spots than MiSeq (mean difference=1,397, CI: 522-2,273, *P* = 8.99*10^−5^, Tukey multiple comparisons of means), but insignificant difference in expected number of blind spots. While the relatively low representation (n=25) of NextSeq 500/TruSeq genomes in our study makes this conclusion tentative (with no bootstrapping, we cannot evaluate a confidence interval), the number of blind spots in its single sampling of 25 genomes is fewer than in any of the 100 bootstraps for MiSeq or HiSeq 2500 workflows prepared with TruSeq library prep (Figure 3A). To evaluate the remaining instrument (HiSeq 2000), we compared it to HiSeq 2500 using workflows prepared with the same library prep kit (Shotgun Nextera). HiSeq 2500 had fewer mean blind spots than HiSeq 2000 (mean difference=8,840, CI: 7,964-9,716, Tukey multiple comparisons of means) and significantly fewer expected number of blind spots (difference between 95% CI: 6,040-11,329) (Figure 3 C,D). By combining the results from these comparisons, we tentatively conclude that among the instruments we evaluated, sequencing with NextSeq leads to the fewest blind spots in the *M. tuberculosis* genome. However, examining more genomes sequenced on NextSeq, and evaluating its performance in combination with additional library prep kits is needed to substantiate this conclusion. We cannot rule out non-additive combinatorial effects between instrument and library prep, which could condition instrument bias on library prep.

#### Coverage bias between library preparation methods

We investigated the number of blind spots between genomes sequenced with the same instrument but prepared with different kits. First, to compare Nextera XT, Shotgun Nextera, and TruSeq library preps, we contrasted blind spots between workflows sequenced with HiSeq2500. Among these three kits, Shotgun Nextera had the fewest mean blind spots (estimated difference between means = 10,903, CI: 10,027-11,779, Tukey multiple comparisons of means) and significantly fewer expected number of blind spots (difference between 95% CI: 8,228-13,987) (Figure 3C). To evaluate the remaining library prep, modified Nextera, we compared blind spots between workflows sequenced with HiSeq2000. After combining these two comparisons, modified Nextera library prep appears to markedly reduce the number of positions impacted by coverage bias in *M. tuberculosis*. HiSeq2000/modified Nextera (the only workflow evaluated with modified Nextera) had the fewest expected blind spots among all workflows (difference between 95 CI: 12,740-20,114 compared to HiSeq 2500/Shotgun Nextera), and fewer mean blind spots than the workflow with same instrument and Shotgun Nextera library prep (estimated difference between means=25,863, CI: 24,987-26,738, Tukey multiple comparisons of means). This gap in blind spots between HiSeq 2000/modified Nextera sequencing workflow and the next best performing sequencing workflow (Figure 3C,D), HiSeq 2500/Shotgun Nextera, is particularly impressive considering that HiSeq 2500 outperforms HiSeq 2000 with the same library preparation, and that Shotgun Nextera was the best performing library prep on HiSeq 2500. Overall, this analysis demonstrates that choice of sequencing workflow can alter the range of blind spots in Illumina WGS studies by tens of thousands of positions.

### Common sequence attributes across library preps and instruments challenge sequencing

Next, we investigated how sequence attributes previously implicated in coverage bias associate with Illumina blind spots in *M. tuberculosis*. Extreme GC content (4), tandem repeats (7), homopolymers (4, 12, 13, 48), and palindromes (10, 16) cause coverage bias for some SBS and SBL technologies, but only repeats and GC content are mentioned by Illumina’s documentation and implicated in SBS chemistry biases (15). After isolating the positions meeting general criteria for each attribute, we took an iterative approach (Materials and Methods) to set thresholds for defining each sequence attribute and classified all positions in the H37Rv genome accordingly (Supplementary Table S7, (30)). For all sequence attributes, blind spots became markedly more abundant as the extremity of each increased (Figure 4 A-C).

**Figure 4.**
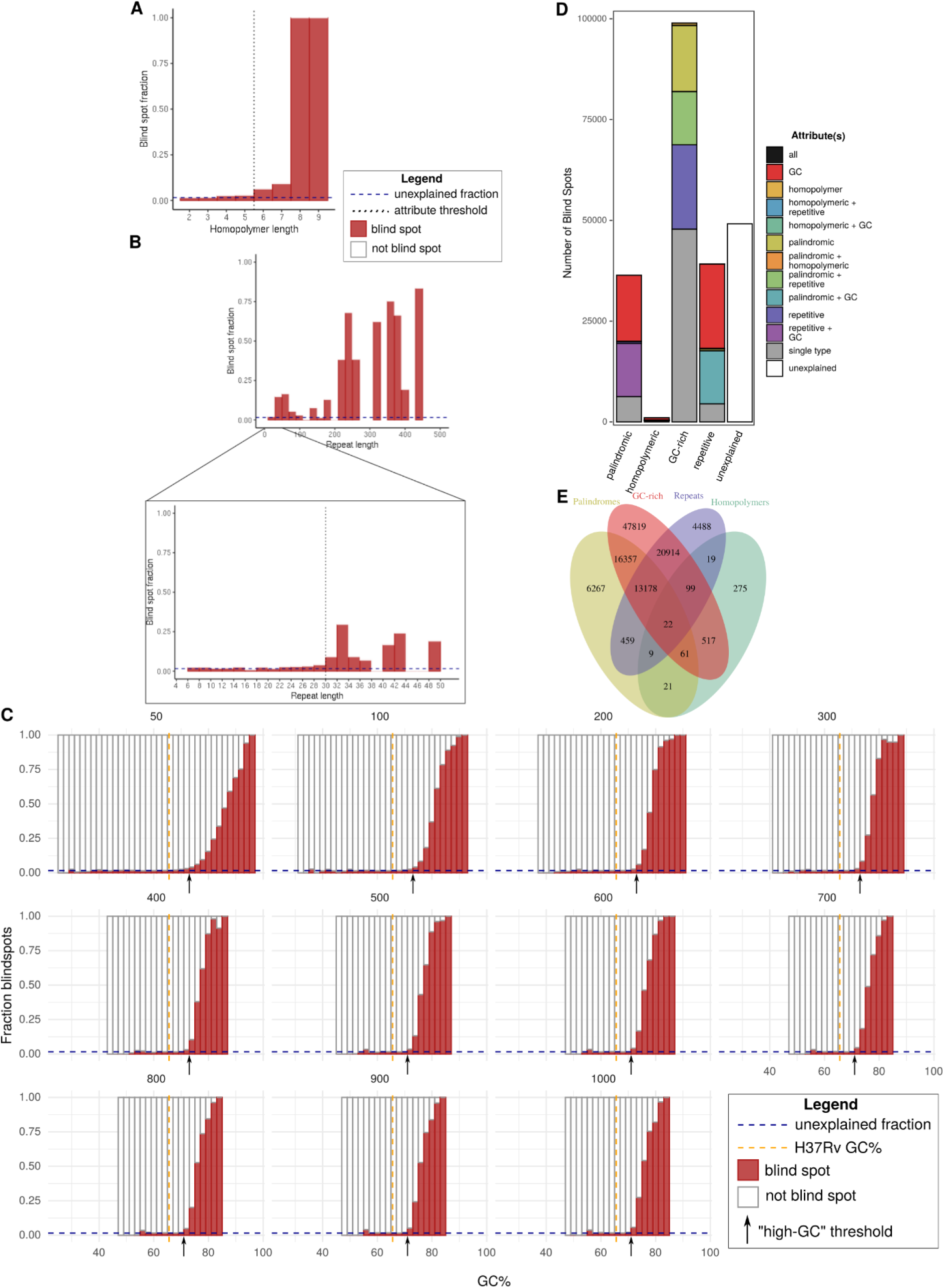
Relationship between blind spot prevalence and sequence attributes previously implicated in coverage bias. **(A-C)** The blind spot fraction (y-axis) among positions binned by attribute-specific parameters. For each attribute, only positions meeting criteria for no other attributes are considered. (**A**) Homopolymers (binned by length, bin size = 1). **(B)** Tandem repeats binned at different lengths to show the general trend between blind spot fraction and repeat length (top, range = 0-500, bin size = 20) and to show precise threshold value (pop-out, range = 0-50, bin size = 2). **(C)** GC-content (window sizes 50 and 100-1,000 by increments of 100). Positions were classified as “high-GC” if they exceeded the threshold for any window size. Palindromes were classified according to previously defined criteria(44). (**D**) The total number of blind spots (y-axis), stratified by sequence attributes (x-axis). Blind spots are grouped and colored according to the set of sequence attributes they meet criteria for. Bar segments are colored as follows: groups of blind spots that only meet criteria for a single sequence attribute (gray), at least two attributes (colors), and those that meet criteria for none of the four sequence attributes (white). (**E**) Number of blind spots meeting criteria for each combination of sequence attributes.

Following classification, we asked how each sequence attribute changed the odds of position being a blind spot. Over half (39,188/75,458, 51.9%) of the bases within 1,588 repetitive regions (≥30 bp) were blind spots, a staggering 37.8-fold (CI: 37.2-38.4) greater frequency than non-repeat regions. GC content was binned by GC% within windows of length 50-1000 (intervals of 100 after 50), and thresholds were calibrated separately for each window size. Because the GC% within the amplified fragment during PCR can dictate coverage bias (4), positions were considered “GC-rich” if they exceeded the threshold in any of the window sizes (Supplementary Table S8). Among 851,580 GC-rich positions, 98,967 (11.6%) were blind spots, 7.58-fold the frequency of non-GC-rich positions (CI: 7.50-7.66). In homopolymeric sequences (length ≥ 6), blind spots were 5.5-fold (CI: 5.28-5.93) more (1,023/5,961, 17.2%) prevalent than in non-homopolymeric sequences. This could be viewed as surprising, considering Illumina maintains that homopolymers do not introduce coverage bias in their SBS technologies(15). Nearly one in seven bases in the *M. tuberculosis* genome (651,837 bases) are within palindromic regions (length ≥ 7), of which 36,374 (5.58%) were blind spots, a 1.7-fold (CI: 1.72-1.76) enrichment compared to non-palindromic sequences, considerably smaller than the other attributes. The modest blind spot enrichment in palindromic sequences also conflicts with previous literature, as they typically only challenge sequencing-by-ligation (SBL) technologies (10). Of the remaining positions that met criteria for none of the four problematic attributes, only 1.6% are blind spots (“unexplained” blind spots).

Next, we asked how being a blind spot changed the odds that a position would possess each sequence attribute, and evaluated overlap between sequencing attributes in the *M. tuberculosis* genome as a potential explanation for the blind spot enrichment in homopolymeric and palindromic sequences. High GC content is considerably more prevalent among blind spots than the other sequencing attributes (62.0% versus 24.5% for repeats, the next highest; Figure 4D). GC content among blind spots (73.0%) is significantly greater (*P* < 2*10^−16^, Fisher’s exact) than the GC content of the H37Rv genome (65.6%) (Figure 4C), and alone “explains” 30.0% (47,819) of blind spots (Figure 4E), dwarfing the number explained exclusively by repeats, homopolymers, or palindromes. Unlike GC-rich blind spots, for other attributes, the overwhelming majority of blind spots are classified as multiple problematic sequence attributes, most often with high GC. Most (69.2%) blind spots can be explained by at least one of these four attributes, yet 49,154 (30.8%) remain unexplained.

While we have described the likelihood of a position being a blind spot given a sequence attribute, we have not done so when attributes are isolated, nor have we evaluated the effect of sequencing workflow on this relationship. To fill these gaps, we calculated the blind spot fraction for each problematic attribute, first by considering all positions meeting the sequence criteria (Figure 5A) and then considering only positions that meet the criteria for one attribute (Figure 5B). Other than for isolated palindromes, the blind spot fraction of all sequencing attributes exceeded the blind spot fraction for unexplained positions, both overall (Figure 5A) and in isolation (Figure 5B).

**Figure 5.**
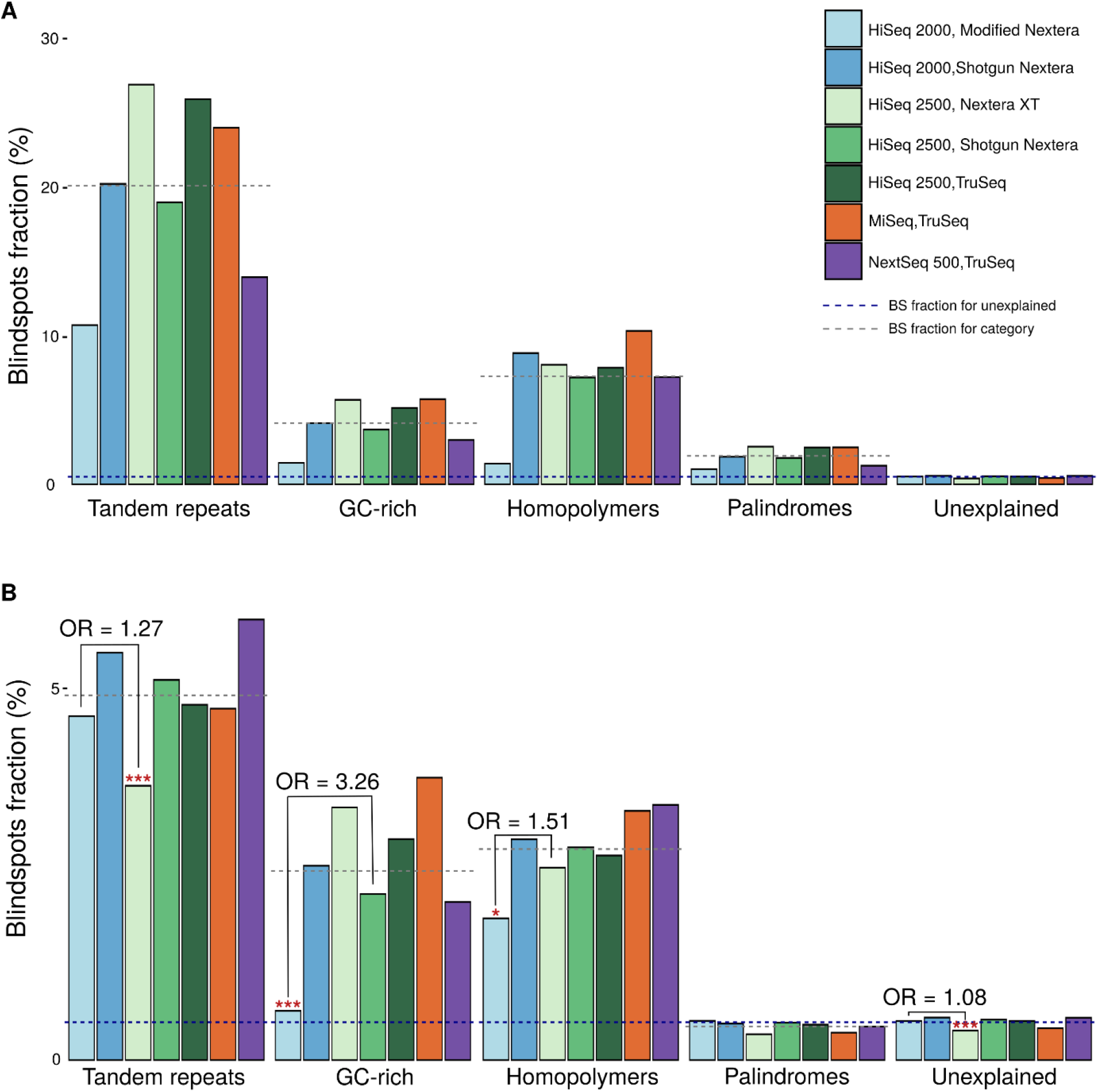
Blind spot prevalence in sequence attributes stratified by instruments and library preps. (**A**) The blind spot fraction (y-axis) in positions with sequence attributes (x-axis) across combinations of Illumina library preps and sequencing instruments (n=175, 25 genomes for each instrument/library prep workflow). Dashed lines denote the blind spot fraction for sequences not meeting criteria for any of the sequence attributes (blue), and the mean blind spot fraction for each attribute across instrument/library prep workflows (gray). (**B**) Same as A, but only considering positions that meet criteria for a single sequence attribute. For each attribute, the workflow with the lowest blind spot fraction was compared to the workflow with the second lowest blind spot fraction, and indicated when their difference was significant (two-tailed Fisher’s Exact, * *P*<0.01, ** *P*<0.001, \*\*\**P*<0.0001, OR=Odds Ratio).

Across all sequencing workflows, tandem repeats are the most problematic attribute, both in combination with other attributes (Figure 5A), and in isolation (Figure 5B). This result is consistent with the literature consensus that repeats introduce mapping ambiguity, creating assembly gaps and depleting coverage (6, 12, 49). It is unsurprising that this problem persists irrespective of library preparation or sequencing instrument, as it is inherent to short-read assembly. While GC-rich regions explain the most blind spots, a given GC-rich position is less likely to be a blind spot than a given homopolymeric or tandem repeat position (Figure 5B). GC-richness accounts for such a large fraction of the blind spots because the entire H37Rv genome is GC-rich (65.6%).The proportion of blind spots in homopolymeric regions was higher than the unexplained blind spot fraction, even in isolation (Figure 5 A,B). This departs from what has been previously described in the literature and refutes Illumina’s documentation that states homopolymers do not cause sequencing errors in Illumina SBS (15).

Isolated palindromic regions lacked coverage bias across all workflows (Figure 5B), consistent with prior reports that palindromic sequences do not challenge SBS methods (10, 18). Curiously, strictly palindromic positions had less coverage bias than unexplained positions (Figure 5B), suggesting additional factors contribute bias and are absent or less prevalent in palindromic regions. This observation may be explained partially or wholly by homopolymers, GC-rich, and repetitive regions with slight coverage bias beyond the resolution of our thresholding criteria (odds ratio >2), and therefore considered unexplained.

### Error profiles differ across sequencing protocols

To investigate blind spot fraction within each attribute, we stratified our analysis across sequencing workflows, using the comparison set of blind spots (Figure 5). By examining differences in coverage bias for the problematic attributes between workflows, we can gain an understanding of the source(s) of coverage bias. The HiSeq 2000/modified Nextera workflow exhibited the least coverage bias in GC-rich (*P* < 2.2*10^−16^; OR=3.26, CI: 3.14-3.37) and homopolymeric (*P* = 0.0091; OR=1.51, CI: 1.10-2.09) sequences, whereas Shotgun Nextera/HiSeq2500 had the least bias (*P*<6.62*10^−8^; OR=1.27, 1.16-1.38) in repeat regions (two-sided Fisher’s Exact test, Figure 5B). HiSeq2000/ modified Nextera also had marginally (OR=1.08, CI: 1.05-1.11) yet significantly (*P*<1.06*10^−09^) less coverage bias in “unexplained” sequences (Figure 5). This could be explained by marginal effects of sequences with GC-rich & shorter homopolymers that show signs of coverage bias (Figure 4 A-C) but did not qualify for these attributes by our definitions.

Across Shotgun Nextera library preps, runs sequenced on HiSeq2000 had more severe coverage bias in all three problematic sequence attributes. This observation makes the comparatively low coverage bias in HiSeq2000/modified Nextera even more remarkable, though it is unclear whether the reductions in coverage bias by modified Nextera (compared to Shotgun Nextera on the same instrument) and HiSeq2500 (compared to HiSeq2000) are additive.

### Common exclusion criteria are neither sensitive nor specific to Illumina coverage bias

While many researchers are unaware of coverage bias, some recognize the problem and address it by omitting large regions of the *M. tuberculosis* genome that are associated with known sequencing biases. These omitted regions are typically restricted to members of the PE, PPE, and PE-PGRS gene families (“PE/PPE genes”), disregarded due to their hypervariable nature, repetitive elements, and propensity for erroneous read mapping (23, 24, 50). However, we identified blind spots beyond these regions. To identify which genes are affected by blind spots, we queried our pooled set of 159,659 blind spots against the H37Rv annotation (NC_000962.3). These blind spots are scattered throughout the genome and overlap 529 genes (Supplementary Table S9). Cumulatively, the PE/PPE genes account for only 55.1% of all blind spots (Table 2, bottom row), meaning almost half of blind spots fall in other regions that are not typically omitted from Illumina sequenced *M. tuberculosis* genomes. Meanwhile, over two-thirds (66.9%) of positions in PE/PPE genes are not blind spots, meaning that many sites within PE/PPE genes are needlessly excluded. Beyond PE/PPE genes, other clinically important genes harbor blind spots, including effectors that subvert human immunity (*esx* genes) (51) and synthesize virulence lipids (*pks* genes) (52), among others (Supplementary Table S9).

**Table 2.**
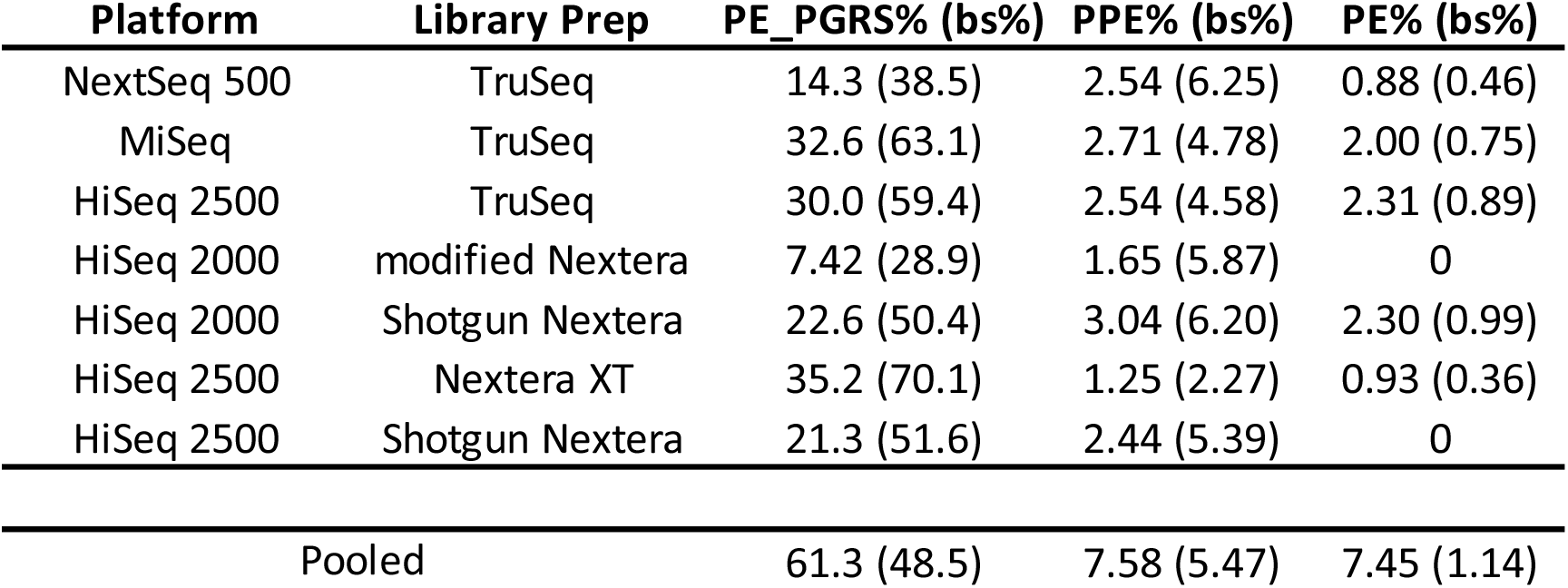
Blind spot prevalence in PE, PPE, and PE_PGRS genes. “PE_PGRS%” **= (**PE_PGRS positions that are blind spots divided by total PE_PGRS positions). “bs%” = (PE_PGRS positions that are blind spots divided by total blind spot positions). Values for PPE and PE columns were calculated analogously (e.g. the blind spot fraction of positions within PE, PPE, and PE-PGRS genes and the fraction of blind spots that fall within each gene set). Fractions were calculated using the “comparison” set of blind spots classified in each workflow (top rows) and the “pooled” set of total blind spots classified within any sequencing workflow (bottom row).

| Platform | Library Prep | PE_PGRS% (bs%) | PPE% (bs%) | PE% (bs%) |
| --- | --- | --- | --- | --- |
| NextSeq 500 | TruSeq | 14.3 (38.5) | 2.54 (6.25) | 0.88 (0.46) |
| MiSeq | TruSeq | 32.6 (63.1) | 2.71 (4.78) | 2.00 (0.75) |
| HiSeq 2500 | TruSeq | 30.0 (59.4) | 2.54 (4.58) | 2.31 (0.89) |
| HiSeq 2000 | modified Nextera | 7.42 (28.9) | 1.65 (5.87) | 0 |
| HiSeq 2000 | Shotgun Nextera | 22.6 (50.4) | 3.04 (6.20) | 2.30 (0.99) |
| HiSeq 2500 | Nextera XT | 35.2 (70.1) | 1.25 (2.27) | 0.93 (0.36) |
| HiSeq 2500 | Shotgun Nextera | 21.3 (51.6) | 2.44 (5.39) | 0 |
| Pooled |  | 61.3 (48.5) | 7.58 (5.47) | 7.45 (1.14) |

Next, we used the comparison set of blind spots (Supplementary Table S5) to investigate which genes are affected by blind spots when using each sequencing workflow. Ninety-two genes contain blind spots regardless of sequencing workflow, while 98 genes contain blind spots only in a single workflow (Supplementary Table S10). HiSeq2000/modified Nextera had the fewest genes affected by blind spots (152 genes) and NextSeq 500/TruSeq had the most (248 genes). We then compared how well the seven sequencing workflows capture the PE/PPE genes. When PE, PPE, and PE-PGRS genes were considered together, the HiSeq 2000/modified Nextera workflow had the fewest blind spots in PE/PPE genes, capturing over 92% of each (sub)family, while the HiSeq 2500/ Nextera XT workflow had more than four times as many and the most among workflows (Table 2).

### Blind spots affect genes implicated in drug resistance

To identify potential blind spots of clinical relevance, we screened the pooled set of blind spots against genes previously implicated in drug resistance (Supplementary Table S3). Eight genes with blind spots have been previously implicated in resistance (Table 3), underscoring the importance of coverage bias in clinical WGS studies. Systematic coverage bias in these genes could obscure resistance signals in Illumina GWA studies, or potentially lead to false associations, if low coverage due to bias were taken to imply deletion. It should be noted that *pks12* may sometimes be lost following prolonged subculturing (53), which would not be filtered out by our phyletic filtering if it arose convergently during culturing. Therefore, some positions in the *pks12* gene (and possibly others) may not reflect true blind spots, but instead selection during culture between isolation and sequencing.

**Table 3.** Resistance-implicated genes with blind spots. Genes implicated is resistance to anti-TB drugs that harbor blind spots. The absolute number of blind spots, proportion of bases in the gene affected (blind fraction), and the PubMed ID of the study that implicated the gene in resistance (PMID).

| gene | # blindspots | blind fraction | drug | PMID |
| --- | --- | --- | --- | --- |
| *pks12* | 3002 | 24.10% | CIP, MDR | 15328105, 29281674 |
| *iniB* | 170 | 11.80% | INH, ETH | 27665704 |
| *Rv2820c* | 45 | 4.90% | ETH | 29358649 |
| *Rv0194* | 73 | 2.04% | STR, TET | 18458127 |
| *mmpL4* | 47 | 1.62% | RIF | 23431276 |
| *ppsA* | 89 | 1.58% | RIF | 23002228 |
| *ppsD* | 2 | 0.04% | RIF | 23002228 |
| *pks2* | 1 | 0.02% | POA | 27759369 |

CIP: ciprofloxacin, ETH: ethambutol, INH: isoniazid, MDR: multi-drug resistance, POA: pyrazanoic acid, RIF: rifampicin, STR: streptomycin, TET: tetracycline

## DISCUSSION

Despite driving the TB pandemic and being among the most frequently sequenced prokaryotes, *M. tuberculosis* coverage bias is poorly understood, and dealt with in varied ways across research groups. Here, we implemented a framework to identify high confidence blind spots from publicly available genomes, stratified by sequencing workflow, demonstrating consistent platform-wide coverage bias in repetitive regions and workflow-specific degrees of difficulty with GC-rich and homopolymeric sequences. We used a stringent false positive rate to capture blind spots with high specificity. As a result, sensitivity is limited by the number of genomes examined. The number of blind spots we identified is a conservative estimate and could be expanded with additional genomes.

Comprising nearly 10% of the genome’s coding capacity and unique to *Mycobacteria*, PE/PPE genes were chief among the interests following publication of H37Rv genome sequence in 1998 (27). However, despite repeated implication of PE/PPE genes in TB pathogenesis, their recalcitrance to Illumina sequencing has led to their near uniform exclusion from many WGS studies. Our results refute the utility of this practice, demonstrating that while many PE/PPE genes contain blind spots (Table 2), numerous others—in some cases entire PE/PPE genes—do not suffer from severe coverage bias with Illumina sequencing. On the other hand, we find that tens of thousands of other positions typically included in Illumina WGS studies harbor positions with severe coverage bias and warrant exclusion (Supplementary Tables S2 and S5). These findings provide more granular criteria for handling Illumina bias in the *M. tuberculosis* WGS.

GC bias is a known issue for Illumina sequencing, yet it is rarely addressed in *M. tuberculosis* WGS studies. Although repeats are the most problematic when present (Figure 5), our results show that 10-fold more blind spots can be attributed exclusively to high GC content than exclusively to repeats (Figure 4E). While increasing the amount of native DNA can improve coverage across the genome and alleviate the effect of GC bias, it is a common misconception that increasing DNA through amplification can have a similar effect and improve coverage uniformly. Instead, amplification bias will further exacerbate the problem in GC-rich regions (Figure 6). Remarkably, however, the modified Nextera library preparation virtually eliminates blind spots in isolated GC-rich positions (Figure 5B). The superior performance of this protocol (Figure 5) was remarkable, particularly considering it was introduced more as a cost-saving measure than one to mitigate coverage bias (29). The modified Nextera protocol substitutes the standard Nextera DNA polymerase with a high-fidelity DNA polymerase enzyme (29), which mitigates PCR amplification bias (54). We recommend this modified Nextera library prep for short-read sequencing of *M. tuberculosis*, and encourage further development of methods that reduce amplification bias.

**Figure 6.**
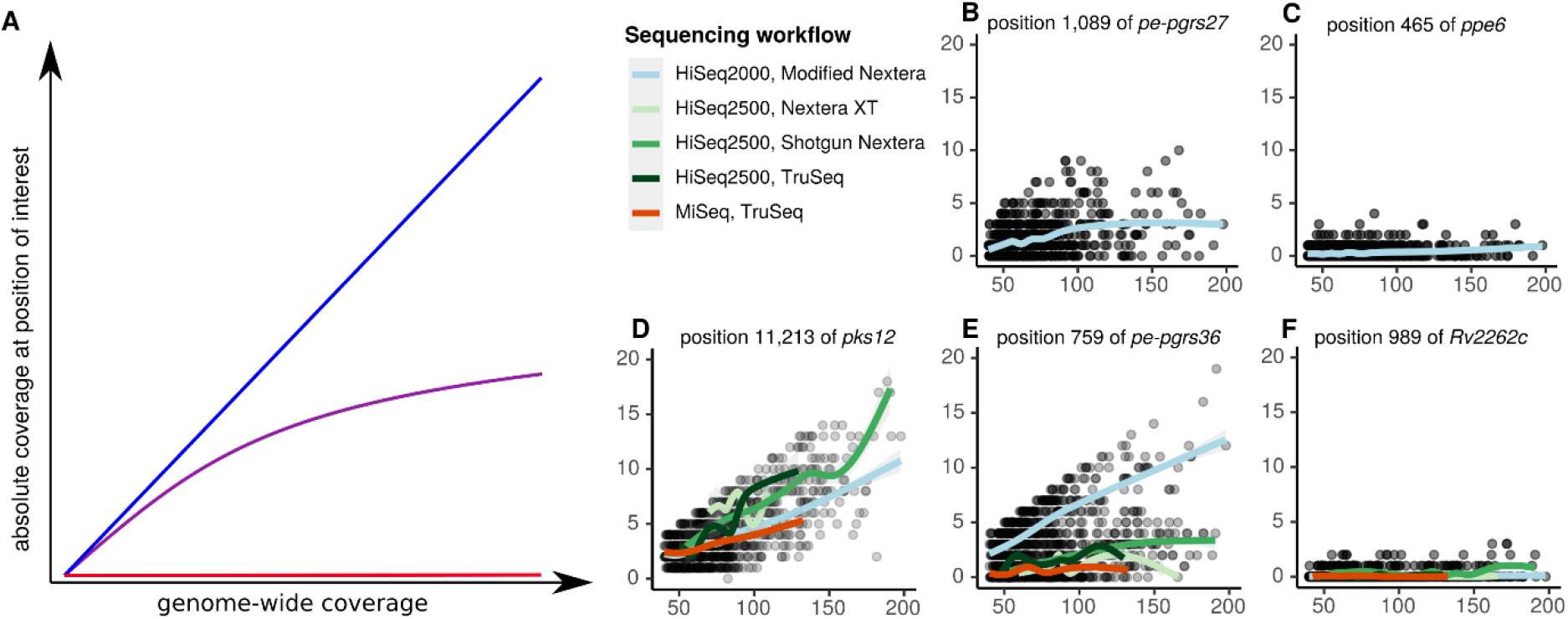
Relationship between mean coverage across the genome and coverage depth in blind spots. (**A**) Illustrations of theoretical linear (blue), sublinear (purple), and constant (red) functions of dependence between coverage depth in blind spots and genome-wide coverage. The type of function relating coverage depth in a blind spot and mean coverage is determined by (i) the nature and source of the coverage bias and (ii) the method for increasing mean coverage. At positions with bias resulting from events with fixed likelihood, coverage depth would increase linearly (blue) with mean coverage, with slope equal to the frequency of the event biasing coverage. For instance, a position situated a number of base pairs away from a large repetitive element, such that 5% of the time the position ends up on the same read as the repeat, would fail to map on those occasions. Therefore, the coverage at the position would be 5% of the mean coverage, regardless of its magnitude (i.e. a linear relationship). Sublinear relationships (purple) can form when the source of bias compounds as mean depth increases. This could occur for positions in GC-rich regions during PCR, as the DNA fragments for which the polymerase has greater affinity will be preferentially amplified, increasing their relative abundance and thus the magnitude of bias for the next amplification cycle. The final relationship is a constant function (red), where the absolute coverage depth at the position of interest does not change as mean coverage changes. This can occur when large repeats are unable to be mapped onto the reference genome unambiguously, as mappability does not depend on coverage. Which of the functions depicted in **A** predominates depends not only on the source of bias, but also the method used for increasing coverage. For instance, GC-rich sequences will likely relate sublinearly to average depth if PCR amplification is employed to increase mean coverage, whereas additional growth to generate more DNA would presumably increase linearly (assuming no additional attributes that would challenge mapping are present). (**B-F**) Observed relationships at representative blind spots in our study. While modified Nextera outperformed other workflows overall, there are still many positions where coverage follows a sublinear (**B**) or constant (**C**) function. Conversely, there are also positions that increase linearly with mean coverage across all workflows (**D**). Improved recoverability with modified Nextera is apparent at many high GC positions (exemplified in **E**), ostensibly due to decreased amplification bias of the high-fidelity DNA polymerase used in the PCR step of the modified Nextera protocol. Lastly, there still remain unresolved sources of bias. The position in hypothetical gene Rv2262c (**F**) met criteria for none of the sequencing attributes we examined, yet demonstrates a constant function across all sequencing workflows.

Importantly, average genome-wide coverage seems to affect blind spot coverage differentially (Figure 6). For some blind spots, coverage increases linearly or sublinearly with average genome-wide coverage (Figure 6B), while others remain static (Figure 6 C,F). Depending on the attributes contributing to the bias, some may be salvageable with more amplification (particularly with the modified Nextera library preparation, Figure 6E) while providing more native DNA is required to salvage others. Further experimentation and analysis is required to rigorously define recoverability of each type of blind spot.

Beyond species-specific insights, our findings clarify two contested notions about sequencing attributes contributing to coverage bias: we refute the assertion that homopolymeric sequences pose no issue for Illumina sequencing (15) and find no coverage bias in palindromic sequences. Our finding that homopolymers associate with coverage bias, even in isolation (Figure 5B), is particularly noteworthy considering that *M. tuberculosis* homopolymers are invariably short (≤ 9 bp). Some researchers contend that homopolymers do not drive coverage bias in Illumina SBS (14, 15), and those who support homopolymer-driven coverage bias suggest it is exclusive to long homopolymers (4, 12, 13). Here, we show that even positions within short homopolymers are manifold more often blind spots than non-homopolymeric positions (Figure 4A and 5B). This finding calls into question the prevailing notion that short homopolymers do not bias coverage (15, 48), suggesting instead that they systematically reduce coverage at lengths as short as 6 bp, and perhaps lower (Figure 4A). Though the persistence of this bias in the absence of other known attributes strengthens our conviction that the bias is genuinely homopolymer-driven, we cannot rule out flanking elements contributing to the bias (4).

The observation that the modified Nextera library preparation attenuates homopolymer bias (Figure 5) suggests the bias originates during PCR amplification, presumably by reducing polymerase slippage (55). Unlike GC-rich and homopolymeric positions, modified Nextera did not improve coverage in repeats, consistent with ambiguous mapping driving coverage bias in repetitive elements, an inherent weakness of short-read technologies. Illumina SBS Instruments beyond those examined here are available. However, they do not address the fundamental factors driving coverage bias, but rather improve sequencing affordability and throughput (e.g. NovaSeq and NextSeq 2000), or portability (MiniSeq and iSeq). Long-read technologies (LRT) offer potential solutions for coverage bias, particularly for repeat regions (4). While LRTs are still catching up to the affordability of Illumina short reads, they have recently made significant advances and single-read accuracy (Pacific Biosciences SMRT-sequencing) and portability (Oxford Nanopore). Applying this framework to identify blind spots across LRTs is an important next step toward a comprehensive knowledgebase of coverage bias across sequencing technologies.

By applying a phylogeny-aware coverage analysis framework to map Illumina “blind spots” onto the genome of virulent type strain H37Rv, we provide a comprehensive roadmap for setting workflow-specific exclusion criteria for *M. tuberculosis* WGS studies. The “pooled” set we report (30) encompasses blind spots present across any sequencing workflow, useful for interpreting Illumina WGS data where workflow is not specified and for studies analyzing genomes sequenced with a variety of workflows, as is common in large-scale GWA studies (56, 57). The “comparison” set of blind spots is useful for identifying which positions suffer from coverage bias on a workflow-specific basis. These lists both inform design of future Illumina WGS experiments and provide a lens through which existing data can be interpreted. While we applied this framework to *M. tuberculosis*, it can also be used to systematically evaluate coverage bias in other species without requiring expensive, time-consuming generation of new genomic data.

## DATA AVAILABILITY

Code used to analyze the primary data and produce the figures and tables are available on GitLab. (https://gitlab.com/LPCDRP/illumina-blindspots.pub/). Data used in the analysis is included or referenced in the supplementary tables, Supplementary Table S7 is available at Zenodo: https://zenodo.org/record/3701840#.Xma5TaaVtGo

## SUPPLEMENTARY DATA

**Table S1:** Sequence information and metadata for all 1,547 genomes included in the analysis.

**Table S2:** Positions in the H37Rv reference genome (NC_000962.3) included in the “pooled” set of blind spots with the absence (0) and presence (1) of each blind spot indicated for each sequencing workflow.

**Table S3:** All genes implicated in drug resistance that were investigated for overlap with blind spots.

**Table S4:** Number of genomes used to classify blind spots in each of the 7 sequencing workflows for the “pooled” set of blind spots.

**Table S5:** Positions in the H37Rv reference genome (NC_000962.3) included in the “comparison” set of blind spots that uses uniform sample size of genomes (n=25) across workflows. Absence (0) and presence (1) of each blind spot is indicated for each sequencing workflow.

**Table S6:** Mean, standard deviation, and variance for number of blind spots classified in each sequencing workflow (randomly selected 25 genomes, bootstrapped 100 times). *only one set of blind spots for NextSeq500, TruSeq workflow given the limiting number of genomes (25).

**Table S7:** Binary matrix indicating whether each position in the H37Rv reference genome met (1) or did not meet (0) the qualifying criteria for the following attributes: homopolymer (length ≥ 6), repeat (length ≥ 30), palindrome (length ≥ 7), high GC-content (specific to window size, see table S5), and blind spot (“pooled” set). This is available at https://zenodo.org/record/3701840#.Xma5TaaVtGo

**Table S8:** Thresholds determined for GC-content, calculated separately for each window size.

**Table S9:** All coding sequences affected by blind spots (“pooled” set). Sheet 1: separated by individual genes, Sheet 2: grouped by gene family

**Table S10:** Coding sequences affected by blind spots (“comparison” set) stratified by the seven sequencing workflows

## ACKNOWLEDGEMENTS

We thank Afif Elghraoui for initially conceptualizing the project and for instructive discussions throughout the project. We also thank Derek Conkle-Gutierrez and Matthew Onorato for their careful review and thoughtful feedback that helped improve this manuscript.

## Author Contributions

Conceptualization: S.J.M., F.V., S.M.R.B., and T.S.; Data curation: C.M., S.M.R.B.; Pipeline Development: C.R., S.N.M., S.J.M., C.M., and S.M.R.B.; Formal analysis: S.J.M. and C.R.; Writing—original draft: C.R., S.J.M., C.M., and S.N.M. Writing—review and editing, S.J.M., C.R., C.M., and F.V.; Supervision: F.V., S.J.M, and S.M.R.B..

